# Slow Oscillations Track Acute Stroke Injury but Not Functional Recovery

**DOI:** 10.64898/2026.06.18.732981

**Authors:** Jake Lee, Arnav Ajay Jadav, Eric C. Landsness

## Abstract

A central goal in translational stroke research is to identify neurophysiological biomarkers that index injury severity and provide information about subsequent functional outcome. Cortical slow oscillations (SOs; 0.1–1.0 Hz) are suppressed after ischemic stroke and recover over the following days to weeks. Whether SO recovery actually tracks behavioral recovery, and whether pre-stroke network organization relates to outcome, has not been tested within individual animals. Using longitudinal wide-field calcium imaging in Thy1-GCaMP6f mice (n = 25), we tracked ipsilateral and contralateral SO power across baseline, 24 hours, and one week after photothrombotic stroke of the left somatosensory forepaw cortex, classifying animals by the presence (STI+; n = 14) or absence (STI−; n = 11) of secondary thalamic injury. Acute ipsilateral SO power was suppressed and tracked concurrent behavioral deficit (ρ = −0.718, p < 0.001), remaining associated with deficit after adjustment for infarct volume (partial ρ = −0.448, p = 0.025). By one week SO power had partially recovered, yet its recovery was dissociated from forelimb use. Week 1 SO power showed no association with behavior in any region or hemisphere (all |ρ| ≤ 0.074, all p > 0.5), and SO recovery did not differ significantly between STI groups despite STI+ animals remaining more impaired (p = 0.011). Pre-stroke SO laterality was associated with week 1 behavioral outcome, independent of infarct size (ρ = −0.518, p = 0.008; partial ρ = −0.446, p = 0.026). Acute SO suppression therefore indexes injury severity beyond infarct volume, whereas spontaneous recovery of SO power is not a reliable surrogate biomarker of week 1 functional outcome. Pre-stroke interhemispheric SO balance emerged as an exploratory candidate source of prognostic information, identifying pre-injury brain state as a dimension that warrants prospective validation.

## Introduction

Ischemic stroke is a leading cause of acquired neurological disability worldwide, producing not only focal tissue destruction but widespread disruption of cortical network dynamics that extends beyond the infarct (GBD 2016 Neurology Collaborators, 2019; GBD 2019 Stroke Collaborators, 2021; van Meer et al., 2010; Bauer et al., 2014). A central challenge in translational stroke research is to identify neurophysiological biomarkers that both index injury severity and provide information about subsequent functional outcome. Whether one physiological signal can serve both roles is not guaranteed, because the acute processes that index injury severity need not be the ones that determine subsequent functional outcome.

Cortical oscillations meet both of these demands after stroke. Changes in low-frequency power in perilesional and remote regions are hallmarks of the injured brain and reflect widespread network disruption (Zappasodi et al., 2019; Wu et al., 2016; van Wijngaarden et al., 2016). The same oscillatory activity also carries prognostic information: quantitative EEG measures of low-frequency power and interhemispheric power asymmetry index injury severity and predict functional recovery in patients (van Putten and Tavy, 2004; Finnigan et al., 2007; Vatinno et al., 2022; Cassidy et al., 2020).

Slow oscillations (SOs; 0.1–1.0 Hz) are a prominent low-frequency cortical rhythm associated with alternating neuronal up and down states and shaped by thalamocortical and corticocortical circuits (Steriade et al., 1993a, 1993b; Neske, 2016; Mohajerani et al., 2010). Slow oscillatory activity can be quantified as spectral power within the 0.1–1.0 Hz frequency band. Focal stroke suppresses ipsilesional low-frequency activity, particularly in lesional and perilesional cortex, and disruption of afferent input can strongly suppress cortical SOs (Lemieux et al., 2014; Landsness et al., 2025). The depth of this acute suppression is therefore a candidate index of injury severity.

Injury severity, however, is shaped by more than the cortical lesion. Because SOs depend on thalamocortical circuits, they are also sensitive to secondary thalamic injury: retrograde degeneration of the connected ipsilesional thalamus follows cortical infarction, and this secondary pathology has been linked to worse functional outcome, at times independent of infarct volume (Cao et al., 2020; Johnston et al., 2024). The presence or absence of secondary thalamic injury (STI) therefore marks an axis of injury severity, spanning milder to more severe damage.

Acute suppression is only the first phase of the low-frequency response to stroke. Over the following days to weeks, regional low-frequency activity can shift back toward its pre-stroke organization, and experimentally enhancing sleep slow waves can promote plasticity and functional recovery (Cassidy et al., 2020; Facchin et al., 2020; Landsness et al., 2025). Whether recovery of SO power actually tracks behavioral recovery within individual animals, however, has not been tested.

Wide-field calcium imaging (WFCI) makes this within-animal approach possible. WFCI captures spatially resolved cortical activity across both hemispheres simultaneously with high temporal resolution, allowing ipsilateral and contralateral SO power to be compared directly at each timepoint (Silasi et al., 2016; Cramer et al., 2019). In mice expressing a genetically encoded calcium indicator in cortical excitatory neurons (Thy1-GCaMP6f), WFCI follows SO power, interhemispheric balance, and bilateral SO covariation longitudinally within the same animal, from before injury through the recovery period (Dana et al., 2014; Chen et al., 2013; Landsness et al., 2025). Paired with behavioral testing and measures of injury, longitudinal imaging relates SO power, injury severity, and functional outcome within each animal, from a pre-injury baseline onward.

Here we used longitudinal WFCI in a photothrombotic mouse model to track SO power and forelimb behavior across a pre-stroke baseline, 24 hours, and one week after stroke, in animals spanning a range of injury severity defined by STI. We first asked whether acute SO suppression indexes injury severity, tracking concurrent behavioral deficit beyond the structural lesion. We then asked whether the subsequent SO recovery provides information about subsequent functional outcome or proceeds independently of it. Finally, we examined whether pre-stroke interhemispheric SO balance is associated with functional outcome.

## Materials and Methods

### Animals

All animal procedures were approved by the Institutional Animal Care and Use Committee of Washington University in St. Louis (protocol 23-032) and were conducted in accordance with institutional guidelines and the Society for Neuroscience Policies on the Use of Animals and Humans in Neuroscience Research. Adult hemizygous Thy1-GCaMP6f GP5.5 transgenic mice (JAX stock no. 024276) on a C57BL/6J background (n = 25; male, 8–12 months) were used in this study. Animals were housed under a 12-hour light/dark cycle with ad libitum access to food and water. All surgical procedures were performed under isoflurane anesthesia (1.5–2% in oxygen).

### Photothrombotic stroke model

Focal cortical ischemia was induced using the photothrombotic (PT) stroke model as previously described (Kraft et al., 2018; Bowen et al., 2025). Animals received an intraperitoneal injection of Rose Bengal dye (10 mg/kg in sterile saline) and were positioned under a stereotaxic frame. A 523-nm diode-pumped solid-state laser was applied to the skull overlying the sensorimotor cortex for 10 or 15 minutes of illumination, durations selected to produce smaller or larger infarct volumes respectively. The PT model produces a well-demarcated cortical infarct with minimal primary subcortical involvement under the present experimental conditions, reproducible lesion geometry, and a clearly defined ischemic lesion, making it well-suited for longitudinal imaging studies (Kraft et al., 2018).

### Stroke infarct volume quantification

To quantify stroke volume, mice were anesthetized and perfused, and brains were harvested and sectioned at 40 µm on a sliding microtome. Each section containing infarcted tissue was stained with Cresyl Violet. Brightfield images of stained slices were acquired using a Keyence BZ-X800 microscope. The cross-sectional area of infarcted tissue on each section was traced using ImageJ, and areas were summed and multiplied by section thickness to obtain infarct volume in mm³ (Bowen et al., 2025).

### Classification of secondary thalamic injury

Secondary thalamic injury was classified based on GFAP immunohistochemistry at one week post-stroke. Brain sections through the thalamus were immunostained with a primary antibody against GFAP (G3893, Sigma-Aldrich; 1:400) and an Alexa Fluor 594-conjugated secondary antibody (715-585-150, Jackson ImmunoResearch; 1:500), as a marker of reactive astrogliosis associated with delayed secondary thalamic pathology after cortical infarction (Kim et al., 2021; Cao et al., 2020). Animals were classified as STI-positive (STI+; n = 14) or STI-negative (STI−; n = 11) by visual inspection for the presence or absence of elevated ipsilateral thalamic GFAP immunoreactivity, assessed by an experienced observer blinded to behavioral and imaging outcomes. In the present study, STI status was used primarily as a biologically meaningful stratification of overall injury burden rather than as an independent mechanistic variable.

### Wide-field calcium imaging

Longitudinal wide-field calcium imaging was performed through a bilateral skull-intact plexiglass window preparation as previously described (Silasi et al., 2016). For window implantation, mice were anesthetized with isoflurane (induction 3–5%, maintenance 1–2%); the scalp was incised and retracted, and a custom Plexiglas window was adhered to the dorsal cranium using Metabond dental cement, completely covering the surgical site. Five days of post-surgical recovery were allowed before any behavioral or imaging sessions. All imaging sessions were conducted in awake, head-fixed mice without anesthesia, so that the slow oscillatory activity measured reflects endogenous cortical dynamics rather than an anesthetic state. These Thy1-GCaMP6f mice express the genetically encoded calcium indicator GCaMP6f in cortical excitatory neurons under the Thy1 promoter (Dana et al., 2014). Imaging sessions were conducted at three timepoints: pre-stroke baseline, 24 hours post-stroke, and one week post-stroke. Sequential illumination of the dorsal neocortex was performed using four LEDs (470, 525, 605, and 625 nm). GCaMP6f was excited with the 470-nm LED and fluorescence emission collected using a cooled sCMOS camera at an 80-Hz acquisition frame rate (20 Hz per channel) (Landsness et al., 2025). Fluorescence images were preprocessed using a pipeline described in Brier et al. (2019). Briefly, temporal drift due to photobleaching was removed by fitting and regressing a fourth-order polynomial from each pixel’s time series. Hemodynamic correction was then performed by dividing the GCaMP6f fluorescence signal by the simultaneously acquired 525-nm reflectance signal, correcting for absorption artifacts arising from fluctuations in oxygenated- and deoxygenated-hemoglobin. Images were spatially smoothed with a 5×5 Gaussian filter, and a global signal, computed as the mean time trace across all cortical pixels, was regressed from each pixel to remove globally shared variance.

### Regions of interest and SO power extraction

Two bilateral regions of interest were defined on the ipsilateral and contralateral hemispheres. The lesion region was centered on the cortical territory directly encompassed by the photothrombotic infarct, corresponding to the primary sensorimotor cortex. The perilesional (peri) region was centered on the cortical territory immediately adjacent to but outside the lesion, corresponding to the peri-infarct region with preserved tissue but disrupted afferent input. The two regions were drawn with the same number of pixels. Slow oscillation (SO) power was computed as the integrated spectral power in the 0.1–1.0 Hz frequency band using a fast Fourier transform (Landsness et al., 2025), expressed in units of power spectral density (×10⁻⁶). For longitudinal comparisons, SO power at each post-stroke timepoint was normalized to each animal’s own pre-stroke baseline value and expressed as a percentage of baseline (% BL), to account for inter-animal variability in baseline SO power. Regions of interest were anatomically fixed across all imaging sessions. ROI placement was performed by an investigator blinded to behavioral outcome and STI status.

### Laterality index

To quantify interhemispheric asymmetry of slow oscillation power, we computed a laterality index (LI) for each region at each timepoint, analogous to established measures of hemispheric asymmetry and recent analyses of sleep-oscillation laterality after stroke (Seghier, 2008; Simpson et al., 2023). The LI was defined as LI = (ipsilateral SO power − contralateral SO power) / (ipsilateral SO power + contralateral SO power). This normalization yields values ranging from −1 to +1, where positive values indicate ipsilateral-dominant SO power, zero indicates perfectly symmetric bilateral SO power, and negative values indicate contralateral-dominant SO power.

### Bilateral SO covariation

Interhemispheric relationships in neural activity have been widely used to evaluate network organization and functional recovery following stroke (Bauer et al., 2014; van Meer et al., 2010). We therefore used bilateral SO covariation to assess the extent to which slow oscillation power in the ipsilateral and contralateral hemispheres varied together across animals at a given imaging session. Bilateral SO covariation was quantified as the Pearson correlation coefficient between ipsilateral and contralateral SO power values, computed separately for the lesion and peri regions across animals at each imaging session. Higher covariation indicates that animals with relatively high SO power in one hemisphere also tended to exhibit relatively high SO power in the opposite hemisphere, whereas lower covariation indicates a weaker relationship between hemispheres. Because this metric was computed across animals rather than from within-animal temporal signals, it should be interpreted as a measure of hemispheric covariation rather than functional connectivity.

### Behavioral assessment

Forelimb motor function was assessed using the cylinder rearing test at each imaging timepoint (Li et al., 2025). Animals were placed in a transparent acrylic cylinder, and spontaneous forelimb use during vertical exploration was recorded. The behavioral score was computed as an asymmetry index, defined as (% left forepaw use − % right forepaw use) / (% left forepaw use + % right forepaw use) (Kraft et al., 2018). Because the photothrombotic lesion was induced in the left somatosensory forepaw cortex, positive values indicate preferential use of the left, ipsilesional, less-affected forepaw over the right, contralesional, affected forepaw, with higher scores indicating greater impairment. Behavioral testing was performed by an observer blinded to STI status.

### Experimental design and statistical analysis

All statistical analyses were performed in Python (version 3.8.0) using the SciPy and pandas libraries. Group differences in continuous variables were assessed using the Mann-Whitney U test (two-tailed) for between-group comparisons and the Wilcoxon signed-rank test for within-animal paired comparisons. Longitudinal changes across the three timepoints were assessed using Wilcoxon signed-rank tests for pre-specified pairwise comparisons between timepoints. Unless otherwise noted, comparisons of SO power (across timepoints, between hemispheres, and between STI groups) were performed on baseline-normalized values (% BL). Associations between continuous variables were quantified using Spearman’s rank correlation coefficient (ρ), chosen for its robustness to non-normality and outliers. Pearson correlation (r) was used for bilateral SO covariation analyses. To test whether bilateral SO covariation differed between the 24-h and week-1 timepoints, we used Steiger’s test for dependent, non-overlapping correlations (Steiger, 1980), which accounts for shared subjects across timepoints. Between-group differences in Pearson correlation coefficients were assessed using the Fisher z-transformation test. Outlier detection was performed using the Grubbs test (Grubbs, 1969; α = 0.05). Two animals were identified as significant Grubbs outliers on week 1 SO power (231.7% and 201.5% of individual baseline, respectively) and were excluded from analyses involving week 1 SO power (n = 23 for those analyses, e.g., SO recovery trajectories), with the exception of bilateral SO covariation, for which all 25 animals were retained at every timepoint. These animals were retained in all other analyses, including those examining the relationship between earlier-timepoint measures (infarct volume, baseline laterality index, baseline SO power) and week 1 behavioral outcome (n = 25 for those analyses), because their behavioral data were not statistical outliers and only their SO power measurements were affected. Statistical significance was set at α = 0.05. All data are presented as mean ± standard deviation unless otherwise stated.

### AI-assisted analysis

AI co-scientist tools were used to propose candidate analyses, assist with statistical implementation and figure generation, and support retrieval and integration of published literature for this dataset: OpenScientist (Roberts et al., 2026) and Biomni (Huang et al., 2025). Analysis code was developed with assistance from Claude Code (Anthropic). All analyses were directed by the authors, and all code, statistical outputs, figures, and cited literature were independently reviewed and verified by the authors against the underlying data and source publications. The authors selected the analyses and interpretations reported here and take full responsibility for all results and conclusions.

## Results

### Injury severity defines the physiological range

Twenty-five mice completed all imaging sessions, with fourteen (56%) classified as STI+ and eleven (44%) as STI−. STI+ animals exhibited larger cortical infarcts than STI− animals (0.972 ± 0.666 mm³ vs. 0.147 ± 0.153 mm³; U = 151, p < 0.001; Figure 1A), indicating that STI classification distinguished animals with differing structural injury burden. Infarct size was associated with acute behavioral deficit (ρ = 0.636, p < 0.001, n = 25; Supplementary Figure 1A) and remained predictive of week 1 behavioral deficit (ρ = 0.502, p = 0.011, n = 25; Supplementary Figure 1B).

**Figure 1.**
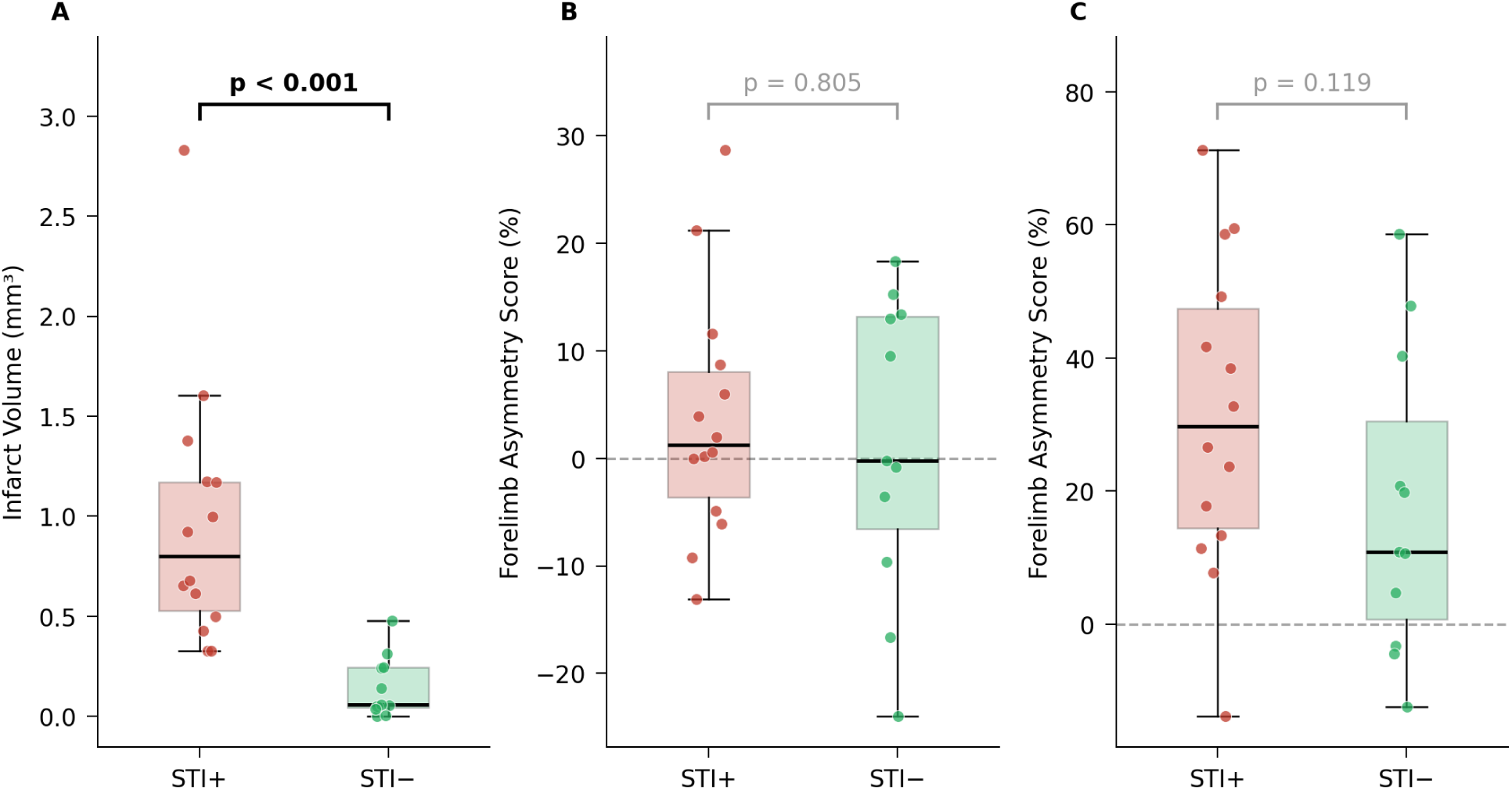
Injury severity defines the physiological range. (A) Infarct volume by STI group. (B) Forelimb asymmetry at pre-stroke baseline. (C) Forelimb asymmetry at 24 h post-stroke (acute). Individual data points shown with box plots (median, IQR, range). STI+ = red, STI− = green. Positive asymmetry values indicate a left forepaw preference, negative values a right forepaw preference.

Baseline behavioral performance did not differ between STI groups (STI+: 3.5 ± 11.4 vs. STI−: 1.3 ± 14.0; U = 82, p = 0.805; Figure 1B). Acute behavioral deficits were numerically greater in STI+ animals, but the difference was not statistically significant (31.3 ± 23.6 vs. 17.6 ± 22.8; U = 106, p = 0.119; Figure 1C), reflecting variability in functional impact despite marked differences in lesion volume.

### Acute slow oscillation suppression indexes injury severity and concurrent behavioral deficit

In the acute phase, SO power was significantly reduced from baseline across both hemispheres and both regions (all Wilcoxon p < 0.001), indicating widespread disruption of cortical slow oscillatory activity. However, suppression was strongly lateralized to the injured hemisphere: ipsilateral SO was significantly lower than contralateral in both regions (lesion: W = 3, p < 0.001; peri: W = 8, p < 0.001). Suppression was also regionally structured, with the perilesional region retaining greater residual activity than the lesion (30.8% vs. 17.2% of baseline; W = 1, p < 0.001), consistent with greater disruption of slow oscillatory activity in the lesion than in the surrounding peri-infarct cortex (Figure 2A).

**Figure 2.**
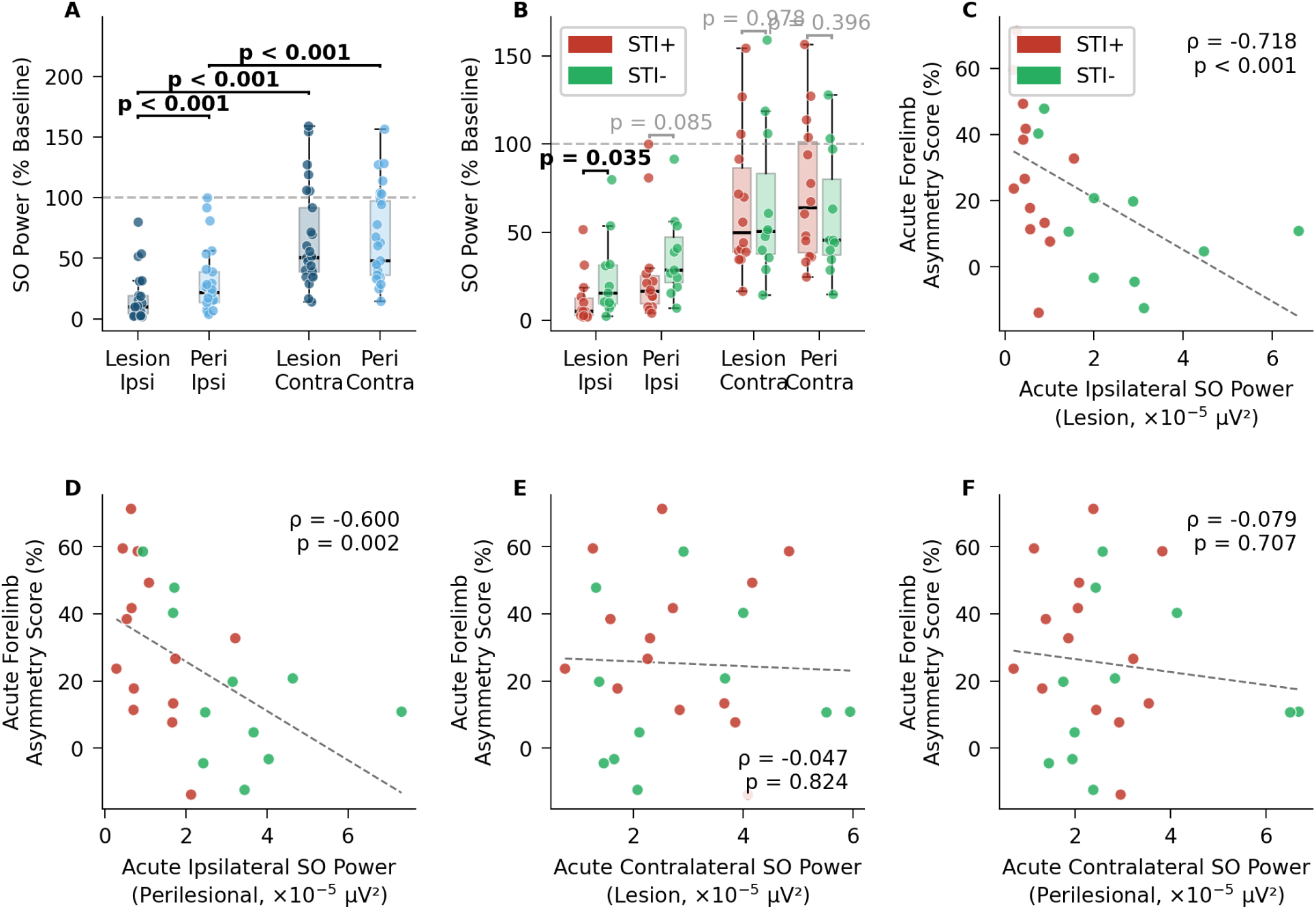
Acute slow oscillation suppression indexes injury severity and concurrent behavioral deficit. (A) Acute SO power across all four region–hemisphere combinations, expressed as a percentage of individual pre-stroke baseline; dashed line denotes baseline (100%). (B) Acute SO power for each region–hemisphere combination stratified by STI group. (C–F) Acute SO power vs. acute behavioral deficit for lesion ipsilateral (C), perilesional ipsilateral (D), lesion contralateral (E), and perilesional contralateral(F). Regression lines shown (C–F). STI+ = red, STI− = green; in panel A, lesion = dark blue, perilesional = light blue.

Ipsilateral SO suppression scaled with injury severity. STI+ animals showed significantly lower ipsilateral SO than STI− animals in the lesion (U = 38, p = 0.035), with a similar but non-significant trend in the perilesional region (U = 45, p = 0.085). In contrast, contralateral SO did not differ between groups in either region (lesion: U = 78, p = 0.978; peri: U = 93, p = 0.396), indicating that while bilateral suppression was observed across the cohort, the magnitude of ipsilateral disruption reflects lesion severity (Figure 2B).

Ipsilateral SO power was strongly associated with concurrent behavioral deficit (lesion: ρ = −0.718, p < 0.001; peri: ρ = −0.600, p = 0.002; n = 25 for both; Figure 2C,D). These relationships remained significant after controlling for infarct size in the lesion (partial ρ = −0.448, p = 0.025), indicating that the association between acute lesion-region SO power and functional impairment remained after adjustment for infarct volume. The perilesional relationship was attenuated after controlling for stroke size (partial ρ = −0.248, p = 0.232), indicating that the perilesional signal is more confounded with injury magnitude. In contrast, contralateral SO showed no association with behavior in either region (lesion: ρ = −0.047, p = 0.824; peri: ρ = −0.079, p = 0.707; Figure 2E,F), confirming that functional relevance is hemisphere-specific. Similar ipsilateral SO-behavior associations were observed within both STI subgroups (STI+: ρ = −0.666, p = 0.009, n = 14; STI−: ρ = −0.673, p = 0.023, n = 11; Supplementary Figure 2).

The laterality index (LI), our measure of the interhemispheric balance of SO power, was likewise strongly associated with concurrent behavioral deficit (lesion: ρ = −0.649, p < 0.001; peri: ρ = −0.612, p < 0.001) and with week 1 behavioral outcome (lesion: ρ = −0.482, p = 0.015; peri: ρ = −0.516, p = 0.008; Supplementary Figure 3A–D). However, acute LI was strongly correlated with infarct size (lesion: ρ = −0.744, p < 0.001; peri: ρ = −0.675, p < 0.001; Supplementary Figure 3E,F), indicating that the acute laterality signal is substantially driven by injury magnitude.

Beyond the lateralized suppression of SO power within each hemisphere, stroke also disrupted the relationship in SO power between hemispheres. Strong baseline SO covariation (lesion: r = 0.865, p < 0.001; peri: r = 0.922, p < 0.001; Figure 3A,B) was disrupted at 24 hours in the lesion (r = 0.213, p = 0.307; Figure 3C), a significant reduction from baseline (Fisher z = 3.64, p < 0.001). Covariation was reduced but remained significant in the perilesional region (r = 0.475, p = 0.016; Figure 3D), also significantly reduced from baseline (Fisher z = 3.60, p < 0.001). This disruption did not differ significantly across STI groups in either region (all within-group acute covariation p > 0.16), indicating that reduced bilateral SO covariation occurred across both STI groups and was not strongly dependent on injury severity.

**Figure 3.**
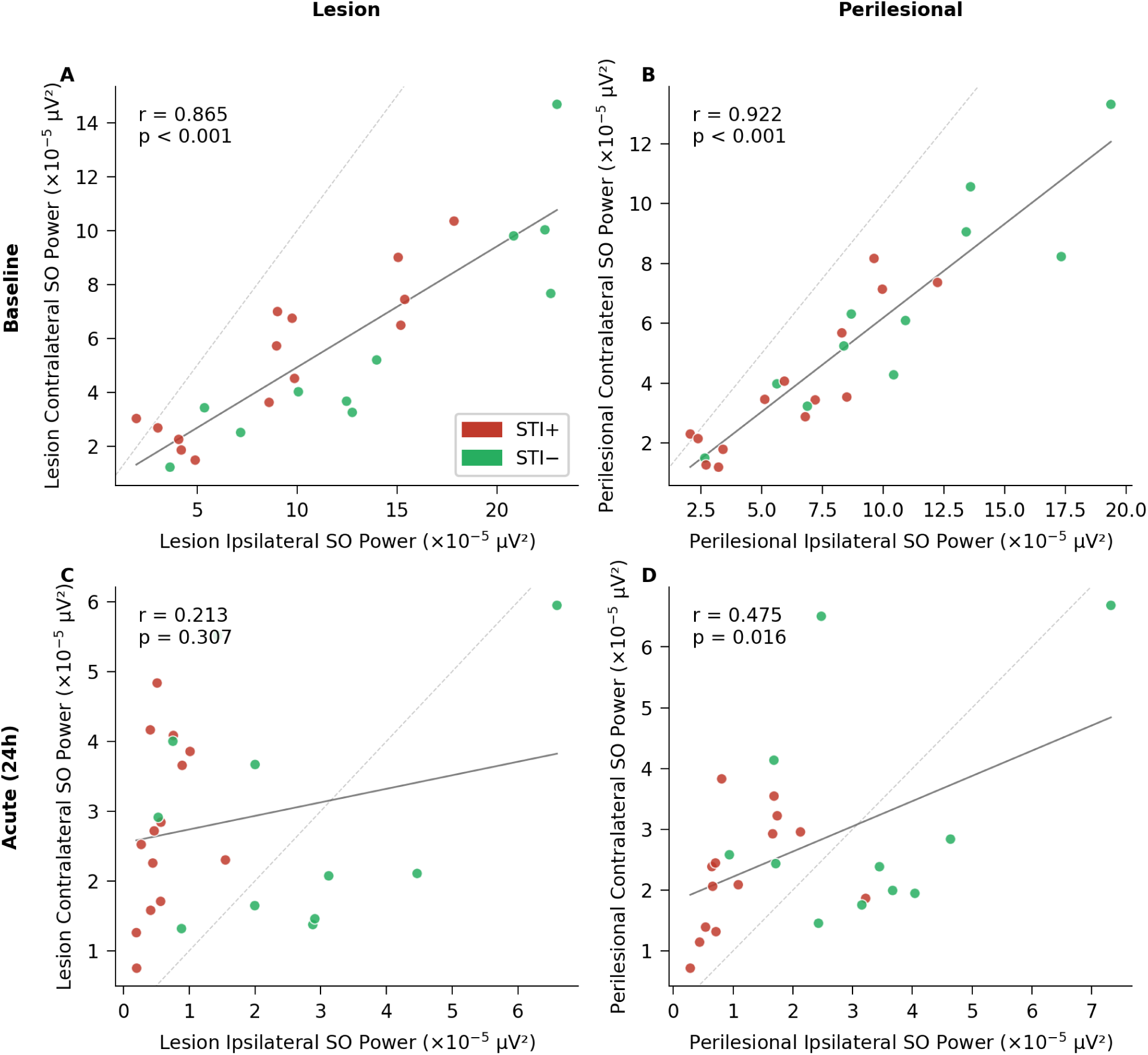
Stroke disrupts bilateral SO covariation. Pearson correlation between ipsilateral and contralateral SO power across animals, at baseline (top row) and 24 hours post-stroke (bottom row), for the lesion (left) and perilesional cortex (right). (A) Baseline lesion. (B) Baseline perilesional. (C) Acute lesion. (D) Acute perilesional. Solid lines are regression fits; dashed lines indicate identity. STI+ = red, STI− = green.

### Subacute physiological recovery dissociates from functional outcome

Having shown that acute SO suppression indexes injury severity, we next asked whether the recovery of SO power over the following week tracks functional recovery, the second question asked of a biomarker. By one week, SO power had shifted toward baseline, but its recovery was incomplete and followed opposite trajectories in the two hemispheres (n = 23, excluding two statistical outliers from trajectory analyses; Figure 4A). Ipsilateral SO, which had been profoundly suppressed acutely, recovered significantly from acute to week 1 in both regions (lesion: 15.0% → 40.3% of baseline, W = 7, p < 0.001; peri: 25.6% → 54.4%, W = 10, p < 0.001). Contralateral SO followed the reverse course: having been only mildly reduced acutely, it did not recover (no significant change in either region) (lesion: 59.0% → 50.2%, W = 81, p = 0.086; peri: 60.3% → 54.6%, W = 106, p = 0.345). The large acute interhemispheric gap thus narrowed through ipsilateral SO recovery rather than contralateral stability. Despite these movements, no region or hemisphere returned to baseline: all four remained significantly below pre-stroke levels at week 1 (all baseline-to-week-1 W ≤ 3, all p < 0.001), indicating that recovery of SO was substantial but partial.

**Figure 4.**
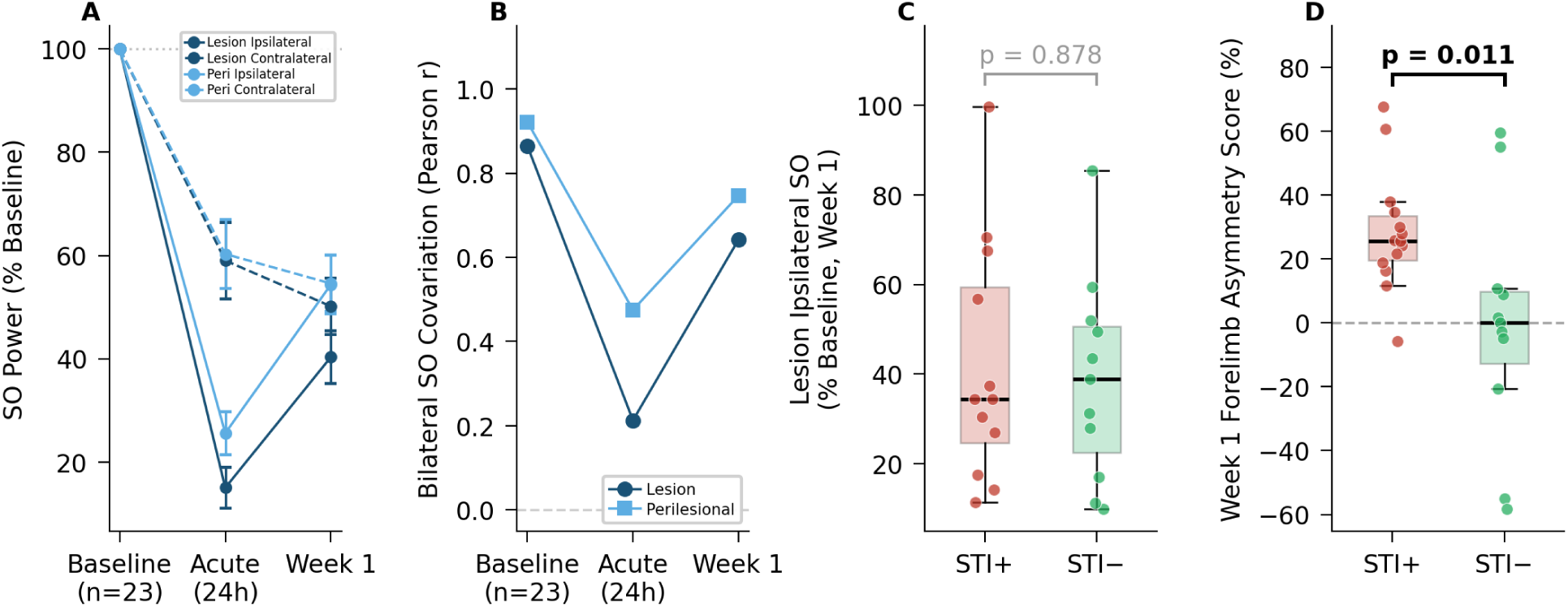
Subacute recovery of SO is partial, regionally divergent, and dissociated from behavioral outcome. (A) SO power across Baseline, Acute (24 h), and Week 1 (% of individual baseline) for the lesion and perilesional regions in both hemispheres (n = 23); dashed line, 100%; error bars, SEM. (B) Bilateral SO covariation across the same timepoints for the lesion and perilesional regions. (C) Week 1 ipsilateral SO recovery (lesion, % baseline) in STI+ vs. STI−. (D) Week 1 forelimb asymmetry in STI+ vs. STI−. STI+ = red, STI− = green; in A and B, lesion = dark blue, perilesional = light blue.

Recovery of SO power also differed between lesion and perilesional regions within the ipsilateral hemisphere. The lesion recovered significantly less than the perilesional region (lesion ipsilateral: 40.3% vs. peri ipsilateral: 54.4% of baseline; W = 24, p < 0.001), and consequently the lesion retained a significant ipsilateral-to-contralateral deficit at week 1 (40.3% vs. 50.2%; W = 52, p = 0.008), whereas no significant hemispheric difference remained in the perilesional region (54.4% vs. 54.6%; W = 120, p = 0.601). The lesion therefore remained the most persistently disrupted region.

Bilateral SO covariation showed a numerical rebound toward baseline. Having been disrupted acutely, covariation rebounded by one week toward baseline levels (lesion: r = 0.642; peri: r = 0.747; n = 25). This acute-to-week-1 increase, however, did not reach significance under Steiger’s test for dependent correlations (lesion: z = −1.71, p = 0.088; peri: z = −1.26, p = 0.207). Across both SO power and bilateral covariation, then, the subacute cortex showed clear signs of recovery of SO toward baseline (Figure 4B).

This recovery of SO, however, did not serve as a reliable surrogate biomarker of functional status. Week 1 SO power showed no association with concurrent behavioral performance across any of the four region–hemisphere combinations (all |ρ| ≤ 0.074, all p > 0.5; Table 1), a null result that held regardless of region, hemisphere, or normalization. The hemisphere closest to baseline at week 1 (contralateral) and the hemisphere that had carried the acute behavioral signal (ipsilateral) were equally uninformative about behavior, indicating that recovery of SO does not track functional state during the first week after stroke.

**Table 1.**
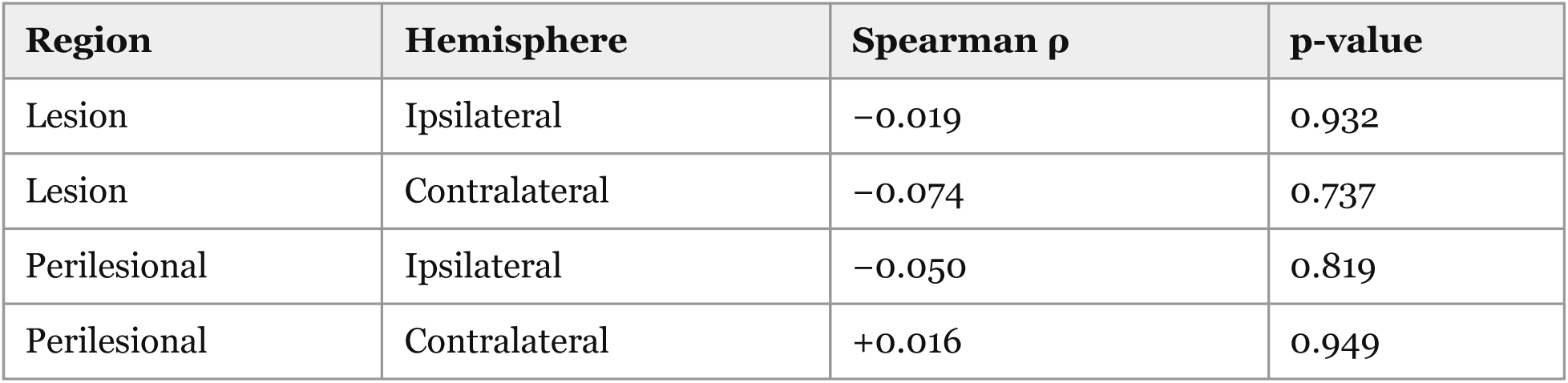
Week 1 SO power is not associated with concurrent behavioral performance. Spearman rank correlations (ρ) between week 1 SO power and week 1 behavioral asymmetry score (n = 23).

| Region | Hemisphere | Spearman $\rho$ | p-value |
| --- | --- | --- | --- |
| Lesion | Ipsilateral | -0.019 | 0.932 |
| Lesion | Contralateral | -0.074 | 0.737 |
| Perilesional | Ipsilateral | -0.050 | 0.819 |
| Perilesional | Contralateral | +0.016 | 0.949 |

The STI comparison made this dissociation explicit. SO recovery did not differ significantly between STI groups: fractional recovery of ipsilateral SO did not differ between groups in any region (lesion: 41.8% vs. 38.8% of baseline; U = 69, p = 0.878; Figure 4C), indicating that SO recovery dynamics were not scaled to injury magnitude. Yet the same animals showed clearly divergent functional outcomes: STI+ animals remained significantly more impaired at one week (28.4 ± 18.6 vs. −0.5 ± 37.0; U = 124, p = 0.011; Figure 4D). Comparable physiology with divergent behavior provides the strongest demonstration that recovery of SO and functional recovery are dissociated over the first post-stroke week.

### Pre-stroke network state is associated with functional recovery

Given the dissociation between post-stroke physiology and functional outcome, we next asked whether pre-stroke network organization determines the capacity for functional recovery. Absolute baseline SO power was not associated with behavioral outcome at one week in any region or hemisphere (lesion ipsilateral: ρ = −0.135, p = 0.519; lesion contralateral: ρ = 0.165, p = 0.432; peri ipsilateral: ρ = −0.188, p = 0.369; peri contralateral: ρ = −0.143, p = 0.495; all n = 25; Supplementary Figure 4). This null result indicates that the overall magnitude of pre-stroke oscillatory activity does not confer an advantage in functional outcome.

In contrast, the baseline LI of the lesion was significantly associated with week 1 functional outcome (ρ = −0.518, p = 0.008, n = 25; Figure 5A). This relationship remained significant after controlling for infarct size (partial ρ = −0.446, p = 0.026), indicating that pre-stroke hemispheric balance provides prognostic information independent of injury severity. Notably, the direction of this relationship was unexpected: animals with greater ipsilateral relative to contralateral SO prior to stroke showed better functional outcomes, a counterintuitive pattern we consider further in the Discussion. Importantly, baseline LI was not significantly correlated with infarct size (ρ = −0.298, p = 0.147; Figure 5B), demonstrating that it does not simply reflect susceptibility to larger lesions but instead may represent a property of baseline network organization associated with functional outcome.

**Figure 5.**
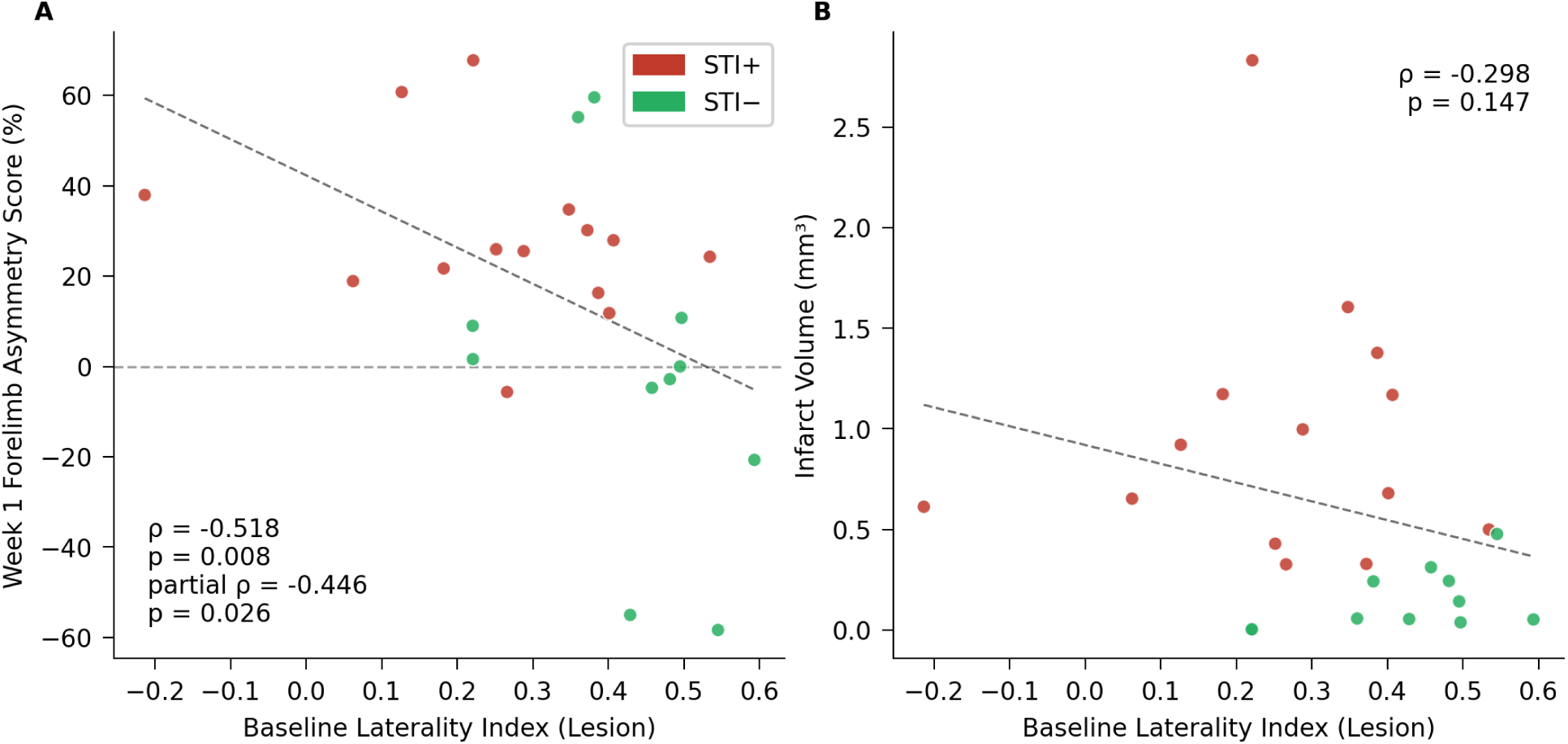
Pre-stroke interhemispheric balance is associated with week 1 functional outcome. (A) Baseline lesion laterality index vs. week 1 behavioral outcome. (B) Baseline lesion LI vs. infarct volume. Regression lines shown. STI+ = red, STI− = green.

A similar but weaker and non-significant trend was observed for the perilesional LI (ρ = −0.021, p = 0.922; partial ρ = −0.092, p = 0.664), suggesting that predictive information is strongest within the lesioned sensorimotor network (Supplementary Figure 5). Together, these findings identify pre-stroke interhemispheric balance as a correlate of functional outcome, suggesting that baseline network organization may contribute information about functional outcome beyond injury severity and post-stroke physiological measures.

## Discussion

We tracked SO power, injury severity, and forelimb behavior within individual animals to ask whether SO power can serve the two roles asked of a stroke biomarker: indexing injury severity and providing information about subsequent functional outcome. Acute SO suppression served the first, indexing injury severity and tracking concurrent behavioral deficit beyond infarct volume. Spontaneous recovery of SO power did not serve the second. Over the first week, SO recovery and functional recovery were not coupled: animals with good and poor outcomes showed comparable recovery of SO. The one physiological measure associated with subsequent functional outcome was present before the stroke rather than after it: the interhemispheric balance of SO power, independent of infarct size.

### Acute SO suppression indexes injury severity beyond lesion volume

Acute ipsilateral SO suppression was graded between the lesion and perilesional cortex, and it tracked concurrent behavioral impairment. The association survived adjustment for infarct volume in the lesion region, indicating that SO suppression captures aspects of functional impairment beyond lesion size alone. The contralateral hemisphere makes the point by contrast: contralateral SO power fell acutely as well, yet had no relation to behavior. The lateralized pattern fits the organization of forelimb control and suggests that ipsilateral SO suppression indexes dysfunction within the affected sensorimotor network rather than a diffuse, whole-brain effect of stroke. This independence from structural damage is what makes acute SO suppression worth considering as a physiological marker of network dysfunction, of a kind that low-frequency EEG measures already approximate in clinical stroke, where quantitative EEG measures such as hemispheric asymmetry and delta-band activity have been associated with neurological severity and subsequent outcome (van Putten and Tavy, 2004; Finnigan et al., 2007).

### Week 1 SO recovery is dissociated from recovery of forelimb use

By one week, the acute coupling between SO power and behavior had dissolved. SO power recovered toward baseline, yet week 1 SO power was unrelated to forelimb use in every region and hemisphere, and bilateral SO covariation followed the same course, indicating that stroke disrupted bilateral SO covariation and not only SO amplitude. The STI comparison makes the dissociation concrete: STI+ and STI− animals showed comparable recovery of SO while remaining unequally impaired at one week. Comparable physiology with divergent behavior provides the strongest demonstration that recovery of SO and functional recovery are dissociated over the first post-stroke week.

The dissociation carries a specific lesson for the use of oscillatory biomarkers after stroke: a signal that recovers need not report that function has recovered with it. These findings suggest that low-frequency physiological measures are most informative when acquired during the acute injury phase, whereas spontaneous recovery of SO power during the first post-stroke week provides limited information about subsequent functional outcome. Although WFCI and EEG measure different physiological signals, both capture low-frequency cortical dynamics and therefore allow related biomarker questions to be addressed. This distinction refines, rather than contradicts, the clinical literature: quantitative EEG measures that predict outcome are typically acquired in the acute or early subacute phase, where they index severity (van Putten and Tavy, 2004; Finnigan et al., 2007; Vatinno et al., 2022), whereas our results concern the recovery trajectory of the same class of signal. Previous work has linked post-stroke low-frequency oscillatory activity to injury state, plasticity, and recovery (Cassidy et al., 2020; Facchin et al., 2020), whereas our within-animal data suggest that spontaneous recovery of SO power can proceed independently of functional outcome during the first post-stroke week. These findings do not argue against oscillation-based therapeutic strategies. Rather, they suggest that spontaneous recovery of SO power should not be assumed to represent functional recovery or serve as a surrogate biomarker of therapeutic efficacy.

Why SO recovery and functional recovery diverge remains an open question. Several processes could plausibly separate them. Homeostatic adjustments in cortical excitability could restore the bulk capacity for synchronized slow activity without reestablishing the specific circuitry that skilled forelimb use requires (Páscoa dos Santos and Verschure, 2022; Fröhlich et al., 2008; Rocha et al., 2024). Reorganization of thalamocortical interactions could return SO power without restoring somatotopic precision (Tennant et al., 2017). Alternatively, recovered slow activity could reflect a cortical state that is less functionally favorable than a return toward normal processing (Sarasso et al., 2025). These accounts are untested here and cannot be distinguished with the present observational design; distinguishing among these possibilities will require direct manipulation of SO activity, a necessary next step toward understanding what recovered SO power represents.

### Pre-stroke interhemispheric balance is associated with functional outcome

The physiological measure most strongly associated with week 1 functional outcome was not any post-stroke measure but a feature of the cortex present before the injury: the interhemispheric balance of SO power. Animals with greater ipsilateral relative to contralateral SO power at baseline had better week 1 functional outcome, and the relationship held after adjustment for infarct volume. Absolute baseline SO power carried no such information, and the effect was confined to the lesioned sensorimotor network, indicating that what mattered was the relational organization of activity between hemispheres rather than its overall magnitude.

Our finding differs in kind from previously described physiological stroke biomarkers because the informative feature was present before the injury occurred and therefore cannot be a consequence of it. Rather than reflecting the brain’s response to stroke, baseline interhemispheric SO balance may reflect a pre-existing property of cortical network organization associated with subsequent functional outcome. This distinction introduces pre-injury brain state as a candidate source of prognostic information that is complementary to conventional post-stroke biomarkers.

We did not anticipate this result, and its direction was itself unexpected. In the cognitive aging literature, pre-existing neural resources are held to buffer the functional consequences of a given amount of pathology, shifting the threshold at which injury produces impairment (Stern et al., 2020). One possible interpretation is that pre-stroke interhemispheric balance represents one manifestation of network reserve, with baseline interhemispheric organization influencing the capacity of surviving tissue to support subsequent functional outcome. This interpretation is broadly consistent with structural and network-based accounts of reserve after stroke (Frontzkowski et al., 2026; Paul et al., 2023), though neither study directly examined pre-stroke SO laterality or ipsilateral dominance. We offer this interpretation as a hypothesis, not a demonstrated mechanism.

Interhemispheric asymmetry is already known to carry prognostic information after stroke: measures such as the brain symmetry index, computed from post-stroke EEG, track severity and predict motor recovery (van Putten and Tavy, 2004; Vatinno et al., 2022). Those measures, however, are obtained after the injury and are therefore shaped by it. Unlike these post-stroke measures, the informative feature in our study was present before the stroke and therefore cannot be a consequence of the injury. The directionality also runs counter to the usual post-stroke pattern, in which greater asymmetry accompanies worse outcome, which underscores that a pre-stroke measure may capture something distinct from its post-stroke counterpart. Given the modest sample and the unexpected direction, we regard the result as exploratory, and one that calls for confirmation in a larger, prospectively designed cohort.

## Limitations

The principal limitation of the study is its observational design: the relationships between SO dynamics and behavior are associative and do not establish causation. Consequently, the mechanisms underlying both the dissociation and the pre-stroke effect remain to be resolved through direct manipulation. Several more specific constraints also apply. STI status was correlated with infarct volume, so the independent contribution of thalamic injury cannot be fully separated from that of cortical lesion size; partial correlations reduce but do not eliminate the confound. The sample size limits statistical power for subgroup comparisons and partial correlations. The findings derive from a single stroke model, cortical target, and behavioral measure, so generalization to other models, lesion locations, and behavioral domains remains to be tested. Recordings extended only through the first post-stroke week; whether the SO–behavior dissociation persists, resolves, or reverses thereafter is a question for longer longitudinal studies.

In summary, SO power indexed acute injury severity, tracking behavioral deficit beyond what lesion volume explained. Over the first post-stroke week, however, SO power recovered without tracking subsequent functional outcome, showing comparable recovery in animals with good and poor outcomes. The physiological measure associated with subsequent functional outcome was present before the injury rather than after it: the interhemispheric balance of SO power, independent of infarct size. More broadly, the results point to the pre-injury state of the brain as an underexplored dimension of stroke prognosis: one accessible today only in preclinical models, but one that may carry prognostic information complementary to conventional post-stroke biomarkers.

## Author Contributions

ECL and JL designed the research. JL and AAJ performed the research and analyzed the data. ECL and JL wrote the manuscript. All authors contributed to manuscript revision and approved the submitted version.

## Generative AI Statement

The authors declare that generative AI was used in the preparation of this manuscript. AI-based tools (ChatGPT, OpenAI; and Claude, Anthropic) were used to assist with language editing. The authors’ use of AI co-scientist tools (OpenScientist; Biomni) for data analysis and figure generation is described in the Materials and Methods. All scientific content, analyses, and interpretations are the authors’ own, and the authors take full responsibility for the content of the manuscript.

## Data Availability Statement

The datasets analyzed for this study are publicly available in WashU Medicine’s Digital Commons Data@Becker Repository (https://doi.org/10.17632/8n5h7v2965). Custom analysis and figure-generation code is publicly available on GitHub (https://github.com/landsness/sti-sleep-oscillations).

---

**Supplementary Figure 1.**
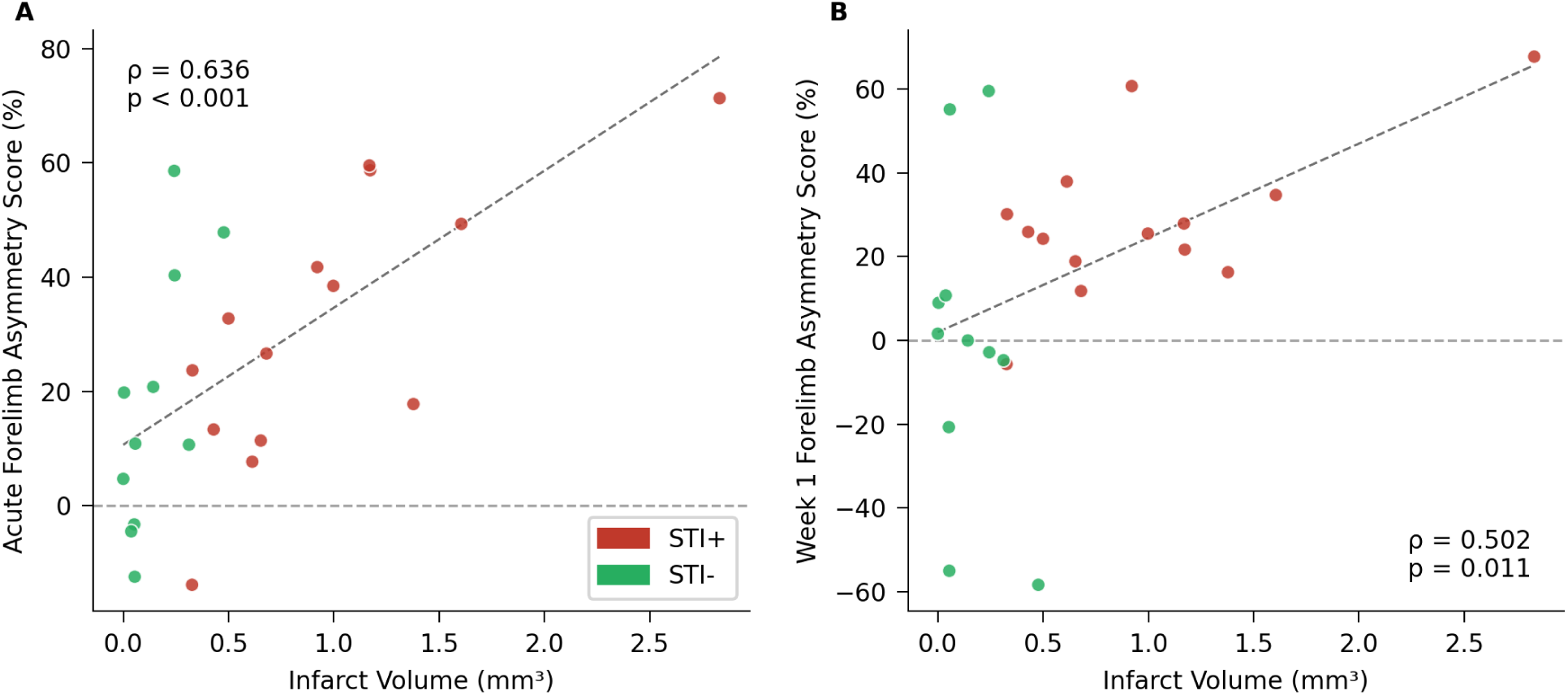
Infarct size and behavioral deficit. (A) Infarct volume vs. acute behavioral deficit. (B) Infarct volume vs. week 1 behavioral deficit. STI+ = red, STI− = green.

**Supplementary Figure 2.**
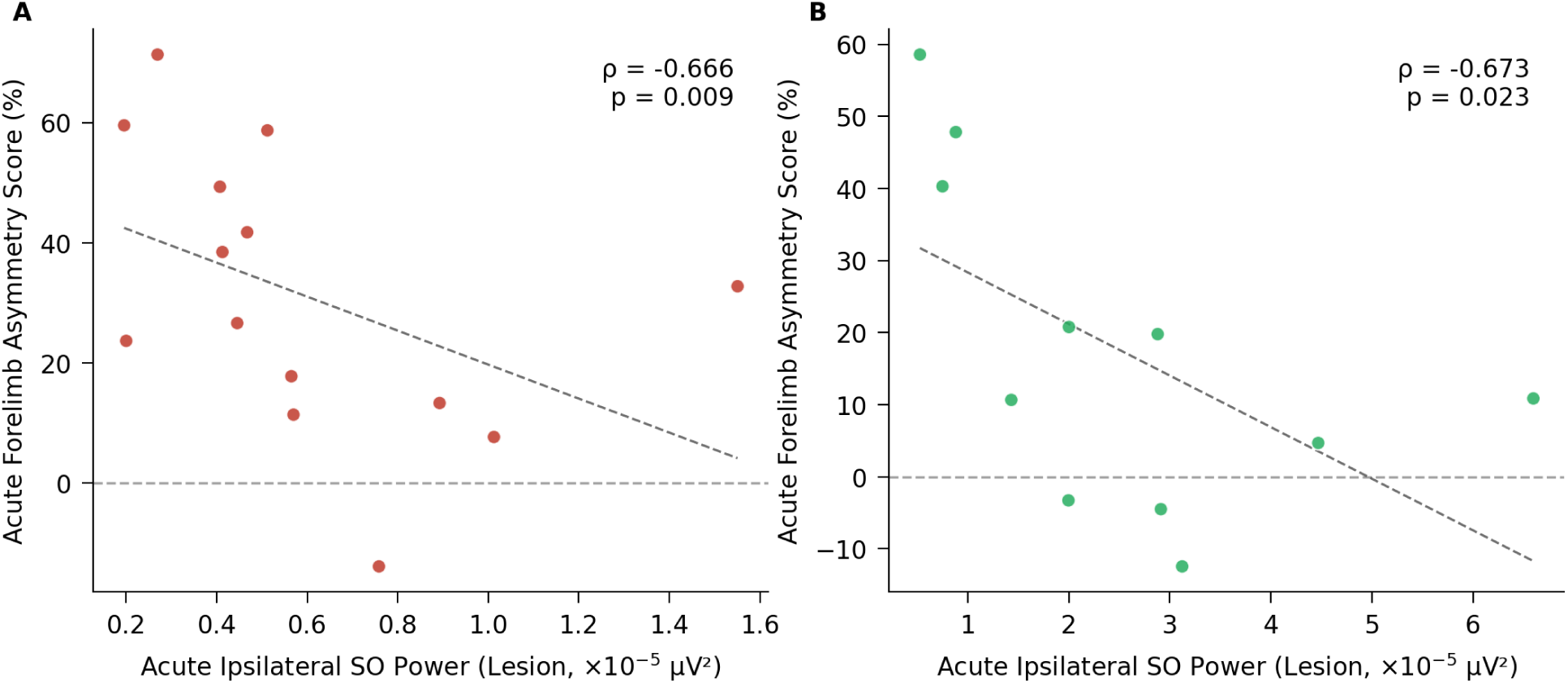
Acute ipsilateral SO–behavior relationship holds within both STI subgroups. (A) STI+ animals (n = 14). (B) STI− animals (n = 11). Acute lesion ipsilateral SO power vs. acute behavioral deficit; regression lines shown.

**Supplementary Figure 3.**
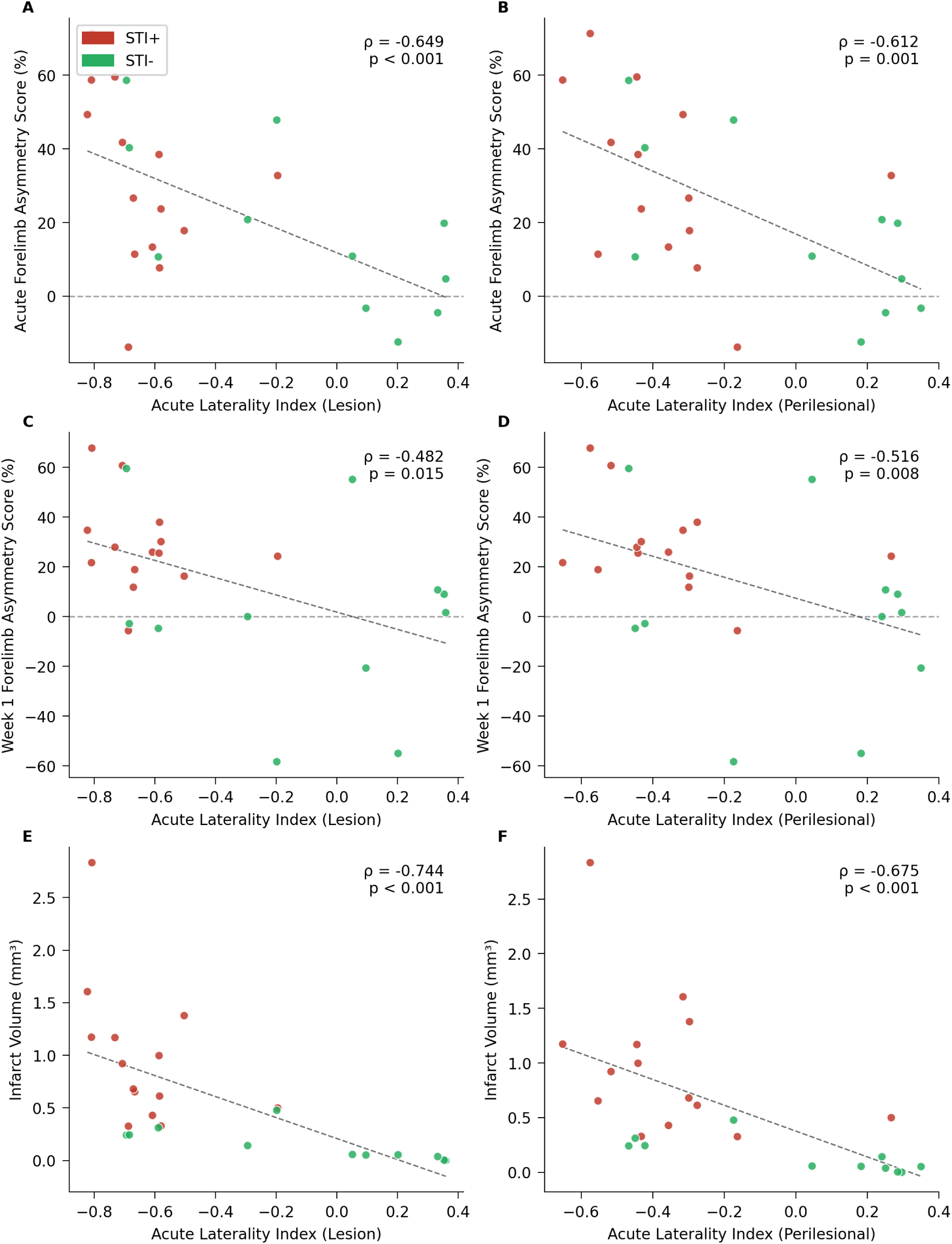
Acute laterality index tracks behavioral deficit but is driven by injury magnitude. (A) Acute lesion LI vs. acute behavioral deficit. (B) Acute perilesional LI vs. acute behavioral deficit. (C) Acute lesion LI vs. week 1 behavioral outcome. (D) Acute perilesional LI vs. week 1 behavioral outcome. (E) Acute lesion LI vs. infarct volume. (F) Acute perilesional LI vs. infarct volume. Regression lines shown. STI+ = red, STI− = green.

**Supplementary Figure 4.**
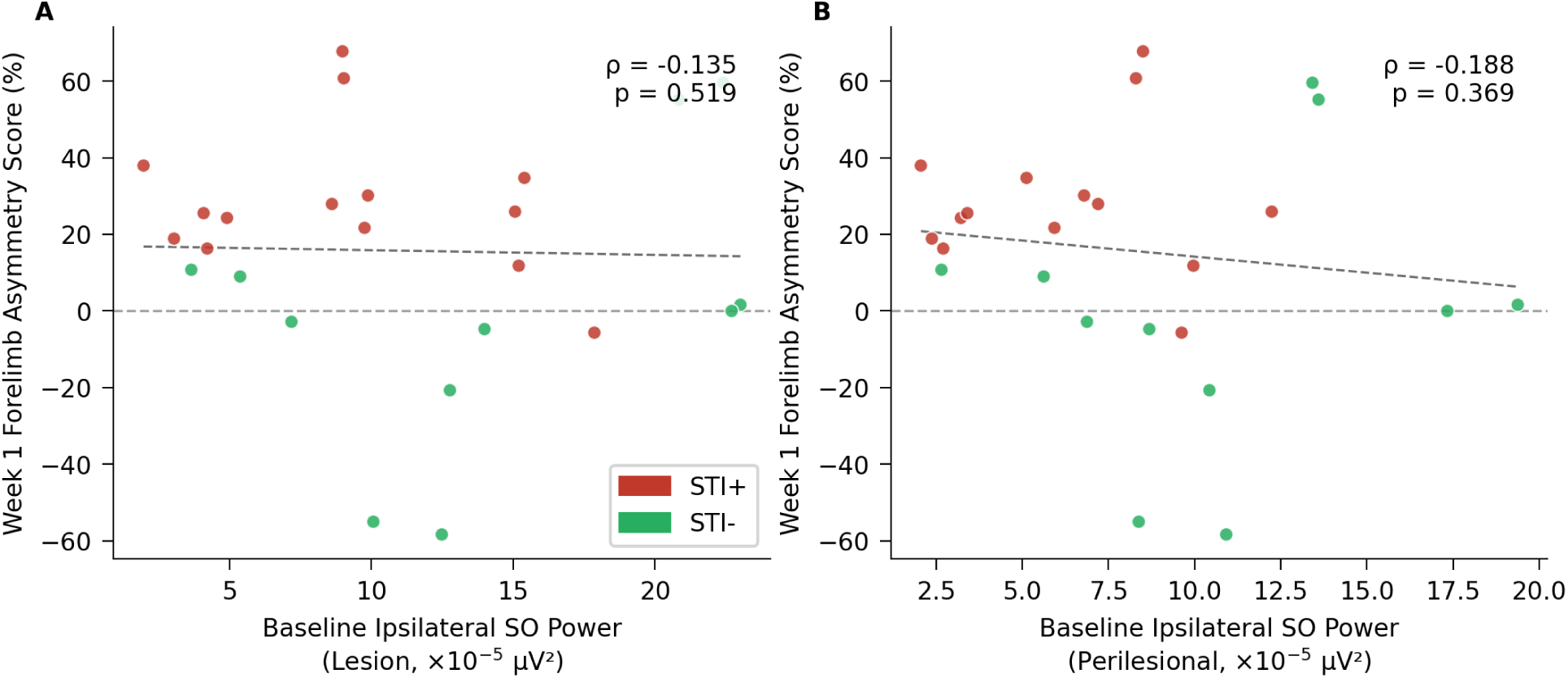
Baseline absolute SO is not associated with week 1 functional outcome. (A) Baseline lesion ipsilateral SO vs. week 1 behavior. (B) Baseline perilesional ipsilateral SO vs. week 1 behavior.

**Supplementary Figure 5.**
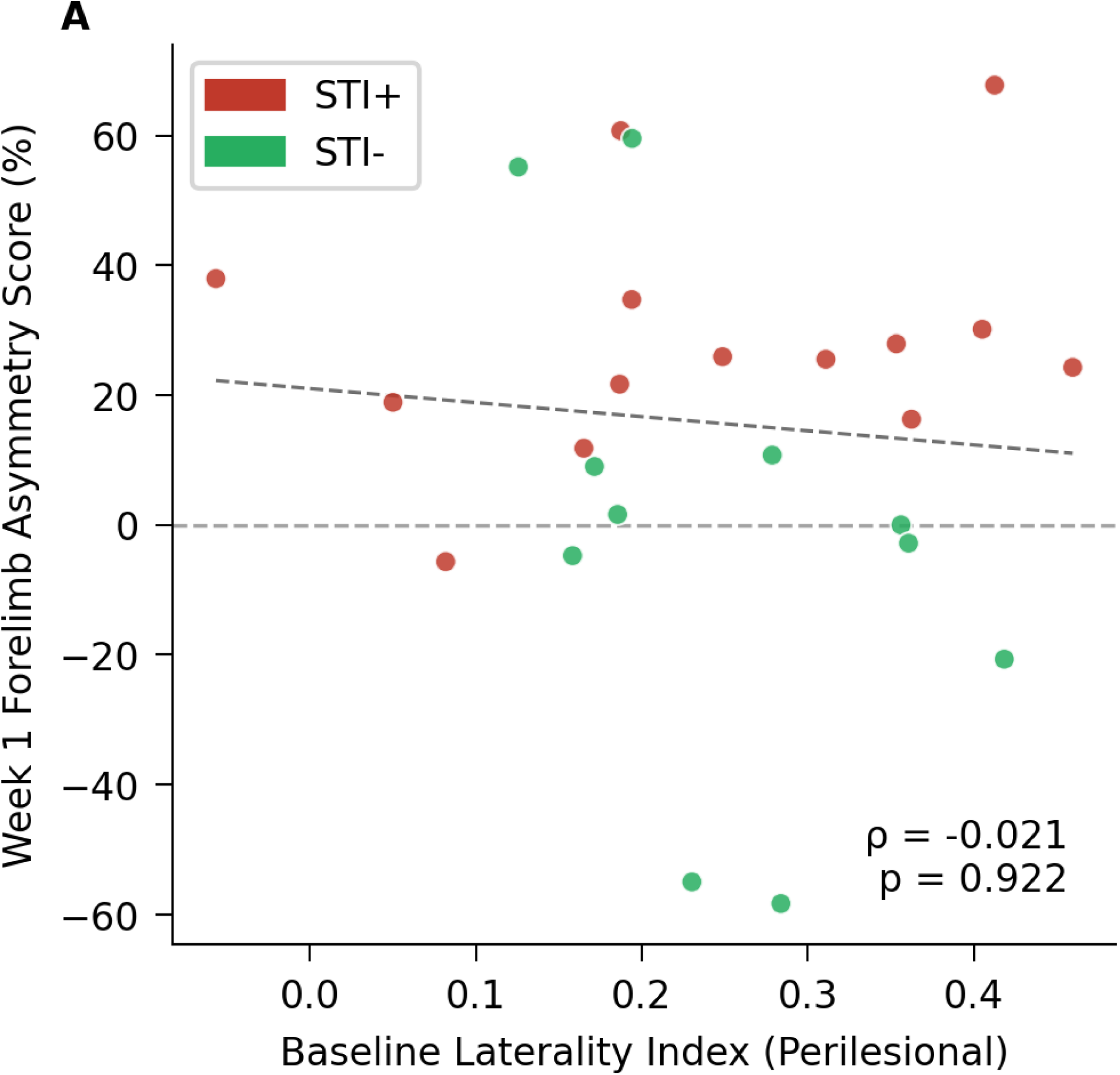
Baseline perilesional laterality index is not associated with week 1 functional outcome. (A) Baseline perilesional LI vs. week 1 behavioral outcome.

## References

1. Bauer, A.Q., Kraft, A.W., Wright, P.W., Snyder, A.Z., Lee, J.M., and Culver, J.P. (2014). Optical imaging of disrupted functional connectivity following ischemic stroke in mice. Neuroimage 99, 388–401. doi: 10.1016/j.neuroimage.2014.05.051

2. Bowen, R.M., Lee, J., Wang, B., Lohse, K.R., Miao, H., Padawer-Curry, J.A., et al. (2025). Early changes in spatiotemporal dynamics of remapped circuits and global networks predict functional recovery after stroke in mice. Neurophotonics 12:S14604. doi: 10.1117/1.NPh.12.S1.S14604

3. Brier, L.M., Landsness, E.C., Snyder, A.Z., Wright, P.W., Baxter, G.A., Bauer, A.Q., et al. (2019). Separability of calcium slow waves and functional connectivity during wake, sleep, and anesthesia. Neurophotonics 6:035002. doi: 10.1117/1.NPh.6.3.035002

4. Cao, Z., Harvey, S.S., Bliss, T.M., Cheng, M.Y., and Steinberg, G.K. (2020). Inflammatory responses in the secondary thalamic injury after cortical ischemic stroke. Front. Neurol. 11:236. doi: 10.3389/fneur.2020.00236

5. Cassidy, J.M., Wodeyar, A., Wu, J., Kaur, K., Masuda, A.K., Srinivasan, R., et al. (2020). Low-frequency oscillations are a biomarker of injury and recovery after stroke. Stroke 51, 1442–1450. doi: 10.1161/STROKEAHA.120.028932

6. Chen, T.W., Wardill, T.J., Sun, Y., Pulver, S.R., Renninger, S.L., Baohan, A., et al. (2013). Ultrasensitive fluorescent proteins for imaging neuronal activity. Nature 499, 295–300. doi: 10.1038/nature12354

7. Cramer, J.V., Gesierich, B., Roth, S., Dichgans, M., Düring, M., and Liesz, A. (2019). In vivo widefield calcium imaging of the mouse cortex for analysis of network connectivity in health and brain disease. Neuroimage 199, 570–584. doi: 10.1016/j.neuroimage.2019.06.014

8. Dana, H., Chen, T.W., Hu, A., Shields, B.C., Guo, C., Looger, L.L., et al. (2014). Thy1-GCaMP6 transgenic mice for neuronal population imaging in vivo. PLoS One 9:e108697. doi: 10.1371/journal.pone.0108697

9. Facchin, L., Schöne, C., Mensen, A., Bandarabadi, M., Pilotto, F., Saxena, S., et al. (2020). Slow waves promote sleep-dependent plasticity and functional recovery after stroke. J. Neurosci. 40, 8637–8651. doi: 10.1523/JNEUROSCI.0373-20.2020

10. Finnigan, S.P., Walsh, M., Rose, S.E., and Chalk, J.B. (2007). Quantitative EEG indices of sub-acute ischaemic stroke correlate with clinical outcomes. Clin. Neurophysiol. 118, 2525–2532. doi: 10.1016/j.clinph.2007.07.021

11. Fröhlich, F., Bazhenov, M., and Sejnowski, T.J. (2008). Pathological effect of homeostatic synaptic scaling on network dynamics in diseases of the cortex. J. Neurosci. 28, 1709–1720. doi: 10.1523/JNEUROSCI.4263-07.2008

12. Frontzkowski, L., Hunze, T.J., Backhaus, W., Bönstrup, M., Gerloff, C., Cheng, B., et al. (2026). The structural reserve of brain networks influences outcomes after a stroke. Brain Commun. 8:fcaf456. doi: 10.1093/braincomms/fcaf456

13. GBD 2016 Neurology Collaborators (2019). Global, regional, and national burden of neurological disorders, 1990-2016: a systematic analysis for the Global Burden of Disease Study 2016. Lancet Neurol. 18, 459–480. doi: 10.1016/S1474-4422(18)30499-X

14. GBD 2019 Stroke Collaborators (2021). Global, regional, and national burden of stroke and its risk factors, 1990-2019: a systematic analysis for the Global Burden of Disease Study 2019. Lancet Neurol. 20, 795–820. doi: 10.1016/S1474-4422(21)00252-0

15. Grubbs, F.E. (1969). Procedures for detecting outlying observations in samples. Technometrics 11, 1–21. doi: 10.1080/00401706.1969.10490657

16. Huang, K., Zhang, S., Wang, H., Qu, Y., Lu, Y., Roohani, Y., et al. (2025). Biomni: a general-purpose biomedical AI agent. bioRxiv [Preprint]. doi: 10.1101/2025.05.30.656746

17. Johnston, P.R., Griffiths, J.D., Rokos, L., McIntosh, A.R., and Meltzer, J.A. (2024). Secondary thalamic dysfunction underlies abnormal large-scale neural dynamics in chronic stroke. Proc. Natl. Acad. Sci. U.S.A. 121:e2409345121. doi: 10.1073/pnas.2409345121

18. Kim, G.S., Stephenson, J.M., Al Mamun, A., Wu, T., Goss, M.G., Min, J.W., et al. (2021). Determining the effect of aging, recovery time, and post-stroke memantine treatment on delayed thalamic gliosis after cortical infarct. Sci. Rep. 11:12613. doi: 10.1038/s41598-021-91998-3

19. Kraft, A.W., Bauer, A.Q., Culver, J.P., and Lee, J.M. (2018). Sensory deprivation after focal ischemia in mice accelerates brain remapping and improves functional recovery through Arc-dependent synaptic plasticity. Sci. Transl. Med. 10:eaag1328. doi: 10.1126/scitranslmed.aag1328

20. Landsness, E.C., Miao, H., Chen, W., Bowen, R.M., Blackwood, S.L., Tang, M.J., et al. (2025). Region-specific non-rapid eye movement delta activity is associated with stroke recovery. Sleep 48:zsaf076. doi: 10.1093/sleep/zsaf076

21. Li, C.X., Kapoor, E., Chen, W., Ward, L.M., Lee, D.D., Titus, A., Reardon, K.M., Lee, J.M., Yuede, C.M., and Landsness, E.C. (2025). Manual assessment of cylinder rearing behavior is more sensitive than automated gait evaluations in young, male mice post-stroke of the forepaw somatosensory cortex. J. Stroke Cerebrovasc. Dis. 34:108325. doi: 10.1016/j.jstrokecerebrovasdis.2025.108325

22. Lemieux, M., Chen, J.Y., Lonjers, P., Bazhenov, M., and Timofeev, I. (2014). The impact of cortical deafferentation on the neocortical slow oscillation. J. Neurosci. 34, 5689–5703. doi: 10.1523/JNEUROSCI.1156-13.2014

23. Mohajerani, M.H., McVea, D.A., Fingas, M., and Murphy, T.H. (2010). Mirrored bilateral slow-wave cortical activity within local circuits revealed by fast bihemispheric voltage-sensitive dye imaging in anesthetized and awake mice. J. Neurosci. 30, 3745–3751. doi: 10.1523/JNEUROSCI.6437-09.2010

24. Neske, G.T. (2016). The slow oscillation in cortical and thalamic networks: mechanisms and functions. Front. Neural Circuits 9:88. doi: 10.3389/fncir.2015.00088

25. Páscoa dos Santos, F., and Verschure, P.F.M.J. (2022). Excitatory-inhibitory homeostasis and diaschisis: tying the local and global scales in the post-stroke cortex. Front. Syst. Neurosci. 15:806544. doi: 10.3389/fnsys.2021.806544

26. Paul, T., Wiemer, V.M., Hensel, L., Cieslak, M., Tscherpel, C., Grefkes, C., et al. (2023). Interhemispheric structural connectivity underlies motor recovery after stroke. Ann. Neurol. 94, 785–797. doi: 10.1002/ana.26737

27. Roberts, K.F., Abrams, Z.B., Cappelletti, L., Moqri, M., Heugel, N., Caufield, J.H., et al. (2026). OpenScientist: evaluating an open agentic AI co-scientist to accelerate biomedical discovery. medRxiv [Preprint]. doi: 10.64898/2026.03.15.26348338

28. Rocha, R.P., Zorzi, M., and Corbetta, M. (2024). Role of homeostatic plasticity in critical brain dynamics following focal stroke lesions. Sci. Rep. 14:31631. doi: 10.1038/s41598-024-80196-6

29. Sarasso, S., D’Ambrosio, S., Russo, S., Bernardelli, L., Hassan, G., Comanducci, A., et al. (2025). Reduction of sleep-like perilesional slow waves and clinical evolution after stroke: a TMS-EEG study. Clin. Neurophysiol. 175:2110746. doi: 10.1016/j.clinph.2025.2110746

30. Seghier, M.L. (2008). Laterality index in functional MRI: methodological issues. Magn. Reson. Imaging 26, 594–601. doi: 10.1016/j.mri.2007.10.010

31. Silasi, G., Xiao, D., Vanni, M.P., Chen, A.C.N., and Murphy, T.H. (2016). Intact skull chronic windows for mesoscopic wide-field imaging in awake mice. J. Neurosci. Methods 267, 141–149. doi: 10.1016/j.jneumeth.2016.04.012

32. Simpson, B.K., Rangwani, R., Abbasi, A., Chung, J.M., Reed, C.M., and Gulati, T. (2023). Disturbed laterality of non-rapid eye movement sleep oscillations in post-stroke human sleep: a pilot study. Front. Neurol. 14:1243575. doi: 10.3389/fneur.2023.1243575

33. Steiger, J.H. (1980). Tests for comparing elements of a correlation matrix. Psychol. Bull. 87, 245–251. doi: 10.1037/0033-2909.87.2.245

34. Steriade, M., Nuñez, A., and Amzica, F. (1993a). A novel slow (< 1 Hz) oscillation of neocortical neurons in vivo: depolarizing and hyperpolarizing components. J. Neurosci. 13, 3252–3265. doi: 10.1523/JNEUROSCI.13-08-03252.1993

35. Steriade, M., McCormick, D.A., and Sejnowski, T.J. (1993b). Thalamocortical oscillations in the sleeping and aroused brain. Science 262, 679–685. doi: 10.1126/science.8235588

36. Stern, Y., Arenaza-Urquijo, E.M., Bartrés-Faz, D., Belleville, S., Cantilon, M., Chetelat, G., et al. (2020). Whitepaper: defining and investigating cognitive reserve, brain reserve, and brain maintenance. Alzheimers Dement. 16, 1305–1311. doi: 10.1016/j.jalz.2018.07.219

37. Tennant, K.A., Taylor, S.L., White, E.R., and Brown, C.E. (2017). Optogenetic rewiring of thalamocortical circuits to restore function in the stroke injured brain. Nat. Commun. 8:15879. doi: 10.1038/ncomms15879

38. van Meer, M.P.A., van der Marel, K., Wang, K., Otte, W.M., El Bouazati, S., Roeling, T.A.P., et al. (2010). Recovery of sensorimotor function after experimental stroke correlates with restoration of resting-state interhemispheric functional connectivity. J. Neurosci. 30, 3964–3972. doi: 10.1523/JNEUROSCI.5709-09.2010

39. van Putten, M.J.A.M., and Tavy, D.L.J. (2004). Continuous quantitative EEG monitoring in hemispheric stroke patients using the brain symmetry index. Stroke 35, 2489–2492. doi: 10.1161/01.STR.0000144649.49861.1d

40. van Wijngaarden, J.B.G., Zucca, R., Finnigan, S., and Verschure, P.F.M.J. (2016). The impact of cortical lesions on thalamo-cortical network dynamics after acute ischaemic stroke: a combined experimental and theoretical study. PLoS Comput. Biol. 12:e1005048. doi: 10.1371/journal.pcbi.1005048

41. Vatinno, A.A., Simpson, A., Ramakrishnan, V., Bonilha, H.S., Bonilha, L., and Seo, N.J. (2022). The prognostic utility of electroencephalography in stroke recovery: a systematic review and meta-analysis. Neurorehabil. Neural Repair 36, 255–268. doi: 10.1177/15459683221078294

42. Wu, J., Srinivasan, R., Burke Quinlan, E., Solodkin, A., Small, S.L., and Cramer, S.C. (2016). Utility of EEG measures of brain function in patients with acute stroke. J. Neurophysiol. 115, 2399–2405. doi: 10.1152/jn.00978.2015

43. Zappasodi, F., Tecchio, F., Marzetti, L., Pizzella, V., Di Lazzaro, V., and Assenza, G. (2019).Longitudinal quantitative electroencephalographic study in mono-hemispheric stroke patients. Neural Regen. Res. 14, 1237–1246. doi: 10.4103/1673-5374.251331

